# The link between gender inequality and the distribution of brain regions’ relative sizes across the lifespan and the world

**DOI:** 10.1101/2025.09.21.677563

**Authors:** Diego Vidaurre, Sergio Morell-Ortega, Pablo Oyarzo, Christopher R. Butler, Marina Charquero-Ballester, Marien Gadea, Floreal Morandat, Boris Mansencal, Pierrick Coupé, José V Manjón

## Abstract

Evidence is emerging that the socioeconomic environment in general, and gender inequality in particular, can be a shaping force on brain structure. However, our understanding of the nature of this influence throughout the lifespan is often limited because most current data sets are geographically and demographically narrow, making it unclear whether results hold across distinct world populations. Here we analyse, for the first time, data from an online MRI analysis platform comprising 13277 subjects from 52 countries and the five continents, across ages that range from childhood to late life. We examined how gender inequality, jointly examined with economic inequality, relates to differences in brain grey matter between males and females. We found that the association between female-male brain differences and gender inequality increases with age, suggesting a cumulative effect of gender inequality throughout life. Further, by considering additional variables that are specifically related to the economy, we found that this effect was, as per current data, dominated by the economic aspects of inequality.

## Introduction

While, at various levels, there is more variability on average within than between sexes (Eliot, Ahmed, Khan, & Patel, 2021; Joel, et al., 2015), machine learning classifiers can still discriminate relatively accurately between male and female brains, even after correcting for brain size (Chekroud, Ward, Rosenberg, & Holmes, 2016). The literature on sex differences often emphasises early development (Ruigrok, et al., 2014), considering two main causes of these differences: the differential make-up of hormones specially during pregnancy and puberty, referred to as the gonadal cause (Young, Goy, & Phoenix, 1964; Arnold & Gorski, 1984); and the fact that all cells in the brain carry either the male chromosome XY or the female chromosome XX, i.e. the genotypal cause (De Vries, et al., 2002; Mallard, et al., 2021). But, at the same time, intertwined with these causes, and accompanying studies in psychology (Guiso, Monte, Sapienza, & Zingales, 2008) and medicine (Mauvais-Jarvis, et al., 2020; Heise, et al., 2019), an emerging literature is unveiling the effect of the environment as a pressing driver of brain differences between the sexes (Baez, Castro-Aldrete, Britton, Ibañez, & Santuccione-Chadha, 2024; Hatzenbuehler, McLaughlin, Weissman, & Cikara, 2024).

Two important recent studies about the link between the brain and structural inequality are (Legaz, Altschuler, …, Bruce, & Ibañez, 2024) and (Zugman, et al., 2023). Legaz et al., examining data from 2,135 older adults, found that economic inequality at the country level (measured with the Gini index) is associated with brain differences across six countries in Latin America and the United States. Zugman et al. analysed 7,876 subjects from different studies, finding a relation between gender inequality and between-sexes brain differences. These two studies were, however, restricted in terms of their age span, with the former focusing on older adults (mean age = 68.6y, s.d. = 8.75y) and the latter on the young (mean age under 25y, maximum age under 35y). Overall, while these and other studies represent progress on the emerging topic of how sociocultural inequality affects the brain, they are either geographically limited or, as mentioned, have limited age ranges. This makes it difficult to assess how gender inequality impacts the relative size of brain regions, beyond brain size, and how it does so through the lifespan in a way that generalises globally.

In this study, we ask the important question of when in the age trajectory gender inequality matters the most for brain structure in a world-representative population, and how it relates to economic wealth and inequality. For this purpose, we analyse, for the first time, a new, large, cross-country dataset from the volBrain web platform. The volBrain platform is a French-Spanish initiative initially developed in 2015 that allows users to perform preprocessing and segmentation of MRI brain scans with minimal effort (Manjón & Coupé, 2016). Specifically, we used data from the vol2Brain pipeline within the platform, which provides a detailed cortical and subcortical segmentation (Manjón, et al., 2022). From vol2Brain, we now have access to 13,335 subjects across the five continents, with ages from childhood to late life. Focusing on grey matter, we found, after correcting for brain size effects, an effect of gender inequality on male-female brain differences in terms of relative grey matter volume across, with a greater effect in subcortical regions. Importantly, we also found that the effect of inequality on male-female brain differences steadily increases with age, suggesting that inequality exerts pressure all through life and not just during development. Our results also suggest a strong economic component underpinning this effect.

## Results

### Demography of the sample

Operating since 2015, the volBrain platform has —at the time of writing— data from over 700,000 subjects from all parts of the world, with ages ranging from infancy to late life. Here, we focus on the vol2Brain pipeline available through the volBrain platform, which gave us access to 13,277 subjects from 55 countries after applying a strict selection criteria (see **Materials and Methods**). **Fig 1A** and **Fig 1B** show basic demographics of the sample, the former in terms of sex and country and the latter in terms of sex and age. Complementing these, **Fig SI-1A** shows the mean age per country in the data set. We gathered, for every country represented in the dataset, two composite metrics of structural inequality: the Gender Inequality Index (GII; from the United Nations’ development report^1^); and the Gini index (from the World Bank’s open data^2^), a classic metric of economic inequality (Ceriani & Verme, 2012). We also gathered the Gross Domestic Product (GDP) as a proxy of average wealth of the countries, which does not consider inequality. The GII, the distribution of which is shown in **Fig 1C** across those countries represented in the vol2Brain dataset, aggregates across three dimensions: health (maternal mortality ratio and adolescence birth rate), empowerment (involvement of women in political leadership and proportion of women with at least secondary education), and labour market participation. The Gini index measures the income or wealth inequality of a country as the departure of the country income/wealth distribution from an ideally egalitarian distribution. Since part of the total income inequality of a country is gender-related (Larraz, 2015), and given the labour market dimension of the GII, these two metrics overlap. This contributes to their high correlation (*r*=0.66; correlations are computed across the 40 countries represented in vol2Brain that have more than 10 subjects) as shown in **Fig 1D**, where the size of the dots represents the number of subjects in the dataset. But they also carry distinct information, because the first two dimensions of the GII are not specific to the economy, and because the Gini index measures economic inequality also between individuals of the same sex. In fact, as observed in the panel, some countries fare better in gender than in income inequality (Switzerland, Italy, Israel, South Africa; in red); and, conversely, some have comparably worse gender equality (Dominican Republic, India, Iran, Egypt and Indonesia; in blue). Furthermore, as also shown in the panel, the GII exhibits a high negative correlation with GDP (*r*=−0.51; correlations are computed across the countries represented in vol2Brain). This anticorrelation is stronger than the one between the Gini index and the GDP (*r*=−0.81; see **Fig SI-1B**) in a statistically significant way (p=0.0005, Meng, Rosenthal and Rubin’s test, (Meng, Rosenthal, & Rubin, 1992))

**Fig 1.**
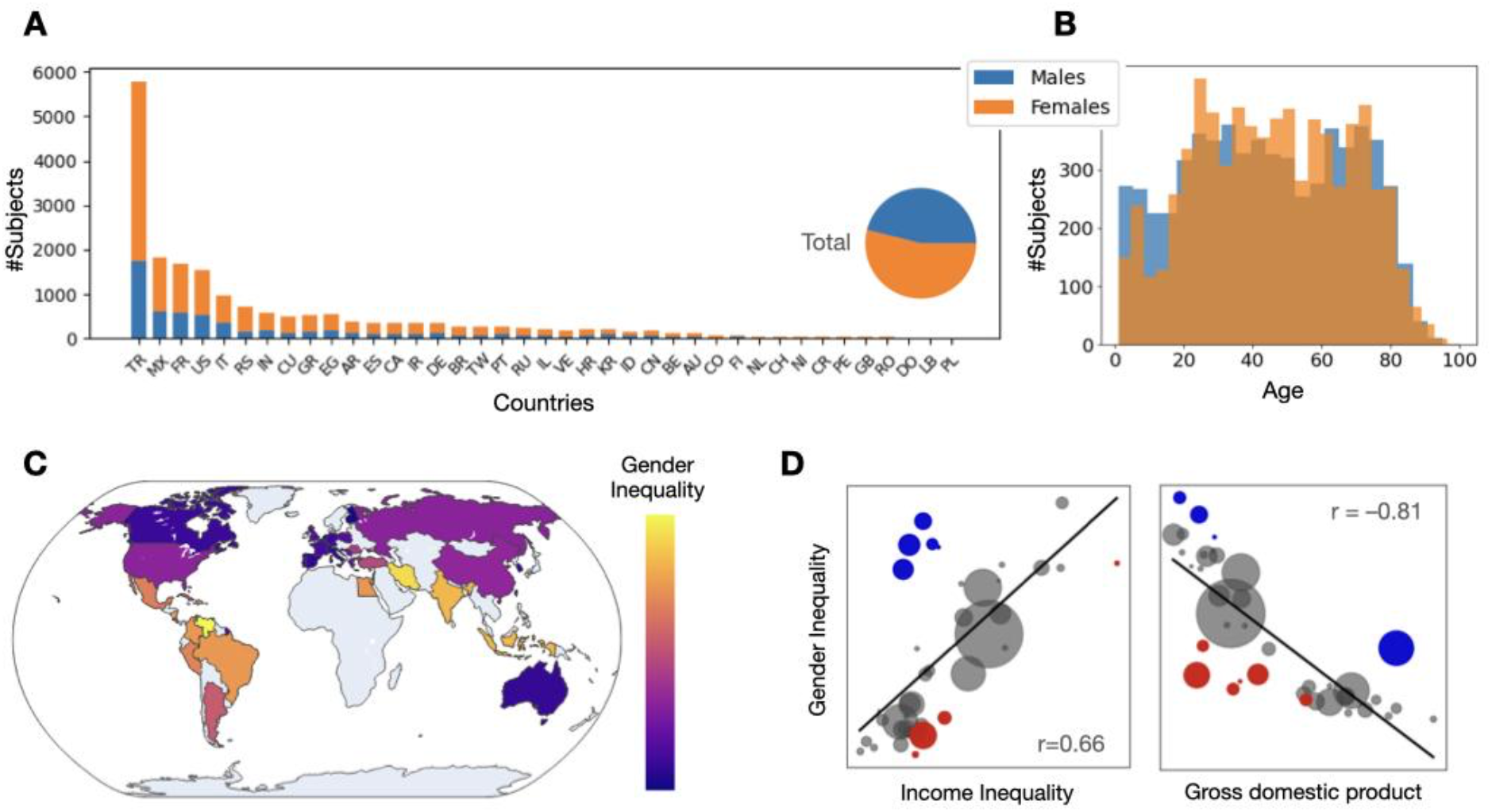
Demographic characteristics of the vol2Brain dataset, from the volBrain MRI processing platform. **A**. Number of subjects per country and sex (for countries with 10 subjects or more), together with the aggregated proportion across countries (pie plot). **B**. Histogram of subjects’ age, per sex. **C**. World map of the Gender Inequality Index (GII) for all countries in vol2Brain. **D**. To the left, scatter plot of income inequality (measured by the Gini index) vs. gender inequality (GII), where the size of the dots represents the number of subjects in vol2Brain dataset from each country; blue indicates countries with relatively worse gender than income inequality (Dominican Republic, India, Iran, Egypt, Indonesia), and red indicates relatively worse income than gender inequality (Switzerland, Italy, Israel, South Africa). To the right, GII vs. gross domestic product (GDP, a proxy of average economic wealth of a country); blue represents countries with relatively worse gender inequality (Dominican Republic, Iran, USA, Venezuela), and red represents countries with relatively weaker GDP (China, Greece, Croatia, S. Korea, Poland, Serbia).

### Relative grey matter volumes across brain regions can predict sex in an age-specific manner

It is controversial whether the differences between female and male brains are not just downstream consequences of brain size. This is because, when properly corrected by brain size, the effects largely reduce (Eliot, Ahmed, Khan, & Patel, 2021). To account for inter-individual differences in overall brain size, we analysed normalized relative volumes. Specifically, grey matter volumes of cortical and subcortical regions (in cm^3^, native space) were scaled such that the sum of all regional volumes equalled 1.0 for each subject. These normalized measures, referred to here as relative volumes, enable a focus on regional distribution patterns independent of total brain volume. Then, using a simple machine learning classifier (based on L_2_-regularised regression) we aimed, in a cross-validated fashion, at predicting sex from the relative region volumes. Leveraging the large number of subjects in the vol2Brain dataset, we ran the prediction per age groups. This was implemented by a 10y-window sliding through an age range from 17y to 80y (i.e. for windows with at least 1000 subjects). Since the windows have different numbers of subjects, we bootstrapped the classifier such that every bootstrapping sample, regardless of the window, had 1000 subjects with a balanced distribution of sex (100 samples). For each bootstrapping sample, and within the cross-validation scheme, we regressed out (deconfounded) intracranial cavity volume (ICV), age (which can vary up to 10y within each window), and the signal-to-noise ratio (SNR) estimation of the image from the brain data (see **Materials and Methods**). Note that ICV is not equivalent to the sum of the region volumes, because ICV accounts for other features of the anatomy such cerebrospinal fluid and white matter. By correcting for both, we took a conservative approach.

**Fig 2A** presents the accuracy curve with a 95% interval of confidence, which was not biased by the number of subjects in each group or the distribution of the classes since these were always equal. Overall, the predictions, while not perfect, are statistically significant and substantially better than chance (i.e. over 0.5).

**Fig 2.**
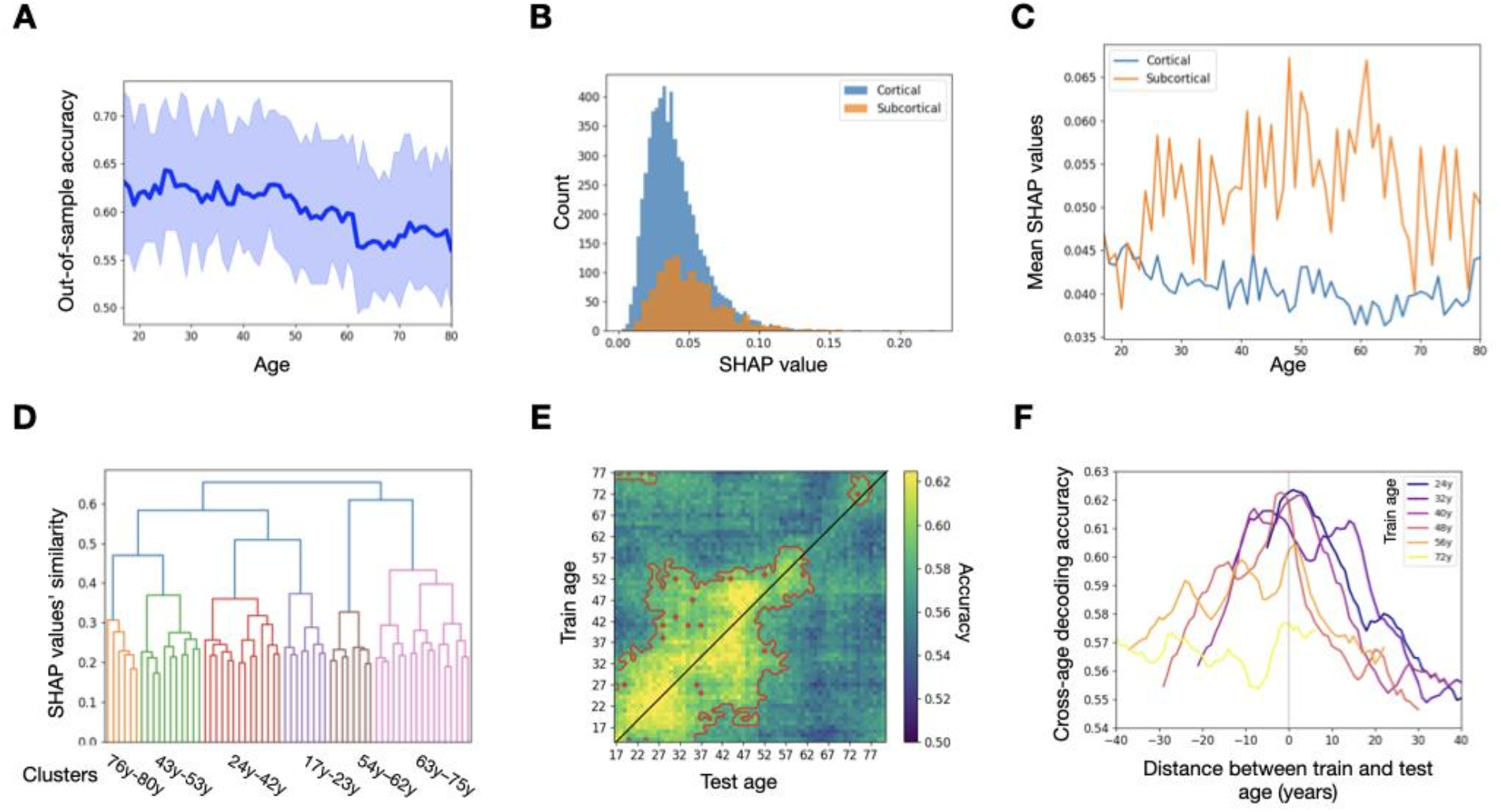
Results from out-of-sample prediction of sex from relative cortical and subcortical grey matter volumes, while correcting for intracranial cavity volume (ICV) and other confounds, indicate that sex can be predicted throughout the life span with relatively good accuracy above and beyond brain size. **A**. Cross-validated accuracy within 10y windows. **B**. Histogram of SHAP values (reflecting the importance of each anatomical feature in the prediction) for cortical and subcortical structures, aggregating across all age windows. **C**. Mean SHAP values for cortical and subcortical regions through the life span. **D**. Clustering of the age windows according to how the anatomical features were used for the prediction of sex, quantified as similarity in their SHAP values. **E**. The age-prediction generalisation matrix, resulting from training a predictive model of sex within every age window and testing on every other age window in a cross-validated fashion. **F**. For a number of example train ages, curves reflecting how the accuracy of sex prediction decreases as the difference between train and test age gets bigger.

To investigate the nature of these predictive models in more detail, we computed, for each age window, the SHAP (Shapley Additive explanations) values associated with the prediction. This metric, inspired by game theory, is increasingly used in explainable machine learning to quantify the importance of each feature in the prediction in a way that takes into account interregional dependencies (Lundberg & Su-In, 2017). **Fig 2B** shows histograms of SHAP values for cortical and subcortical regions across all age windows. **Fig 2C** shows the mean SHAP values (averaged across cortical and subcortical regions separately) as a function of age. Although, as a whole, cortical regions may have more predictive power because there is a larger number of them, one by one the subcortical structures are on average more important for the prediction throughout the age span.

Based on the anatomical patterns of sex prediction as encoded by the SHAP values, we then sought to quantify whether the pattern of anatomical features that underlies sex prediction changes through the life span. We performed a hierarchical clustering analysis where the similarity between two age windows was quantified as the correlation of the SHAP values. **Fig 2D** shows the resulting dendrogram with colours indicating clusters. To further quantify this temporal effect, we set up a cross-validation procedure where we trained a predictive model on subjects of a given age window and tested the model on all the other age windows (always making sure that there is no data leakage between train and test). This produced a (no. windows by no. windows) matrix of cross-validated accuracy that we refer to as the age-prediction *generalisation matrix*. This is shown in **Fig 2E**, where the red contours indicate regions of above-chance prediction (bootstrap-based testing); and in **Fig 2F**, where we show, for a selection of models (each trained on a different age window), how the accuracy declined as test age moves away from train age.

Throughout these results, we can observe that the prediction of sex becomes harder in older age, and that models trained on the younger age ranges do not generalise to older ages and vice versa. Overall, these analyses indicate that the nature of sex differences in the brain changes over time. Next, we investigate possible socioeconomic causes behind this effect.

### Female-male brain differences reflect a cumulative effect of gender inequality across the lifespan

Having established the existence of female-male differences in the sample, we asked if income inequality relates to these differences. For this purpose, using the same 10y sliding windows as in the previous analysis, we computed Euclidean distances between each female-male pair within the window and within each country. For each age window, we then regressed these pairwise distances on the following covariates: the GII of the country where the pair is from, their age difference (which is maximum 10y, i.e. the window width), their difference in ICV, and their difference in SNR. This way, we test whether GII is a significant regressor even in the presence of the other three covariates. We embed the regressions into a cluster-based permutation testing procedure to infer statistical significance of the regressors (Maris & Oostenveld, 2007). To avoid the comparisons between age windows being biased by the fact that the windows vary in the number of subjects, all tests were run on subsamples of 22.000 subject pairs (95% of the number of subject pairs for the 80y-centered window, which contains the lowest number of pairs). To assess the robustness of the estimations, we repeated the analyses five times with different random samples.

**Fig 3A** compares, in terms of explained variance (*r*^*2*^) the full model containing the four covariates (plus an intercept) against a nested model that excludes GII (each sample is represented as a separate curve); the difference, the average of which is represented by the grey area, is significant from early adulthood (p<0.01; the intensity of the pink area reflects how many of the bootstrapped subsamples were significant, with all subsamples turning significant from 28y). This result highlights the importance of GII in explaining female-male grey matter differences. Focusing on the model with GII, **Fig 3B** examines the regression coefficients through the life span. Using the same testing scheme, GII was a significant covariate in all subsamples from 28y onwards, confirming the previous result. The dotted red line indicates the slope of the GII curve, which is significantly positive (t-test, p<10^-18^ for all five repetitions of the analysis), suggesting a cumulative effect of gender inequality throughout life.

**Fig 3.**
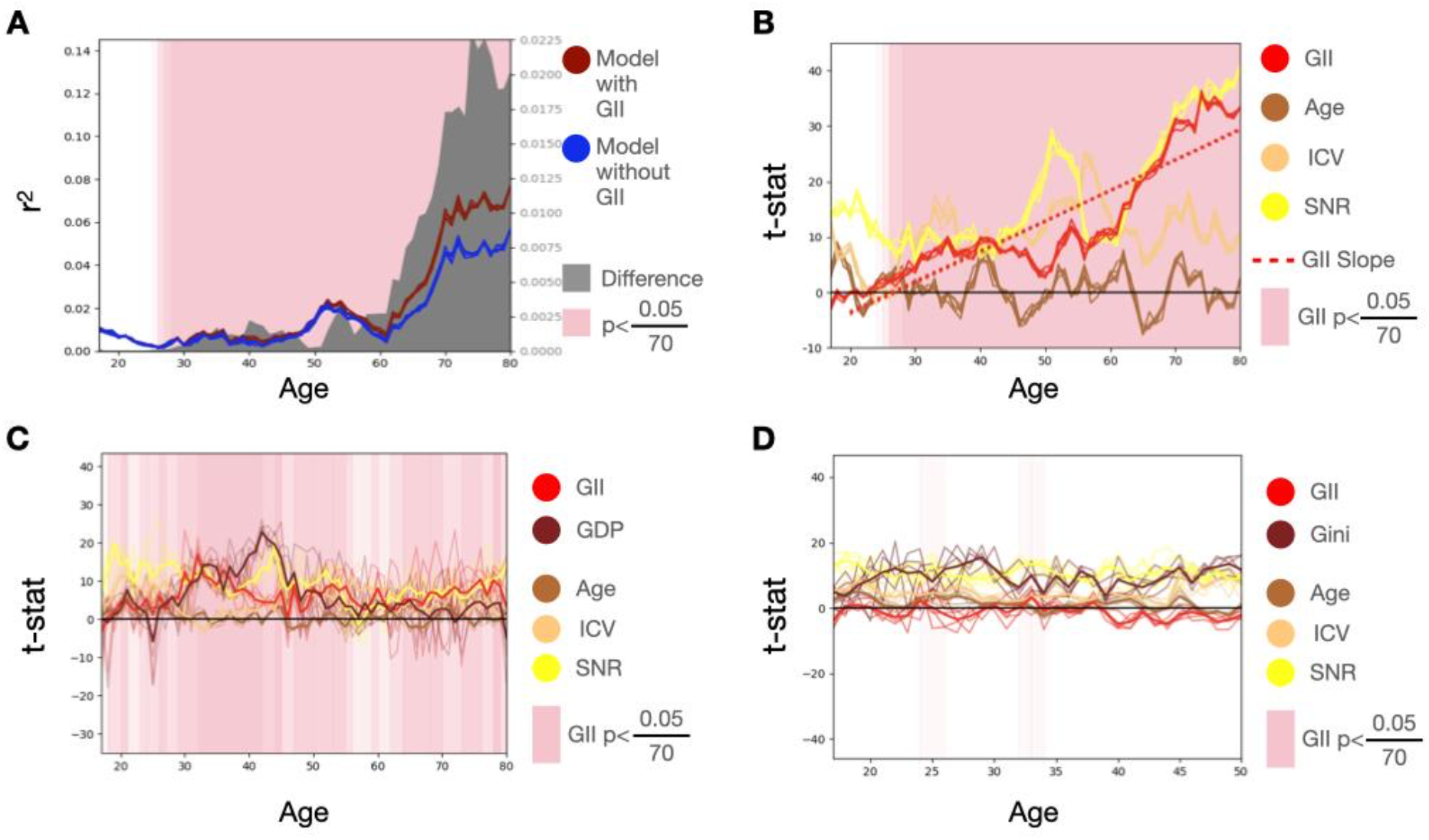
Statistical testing shows a significant relation between-sexes brain differences (corrected for overall brain size by normalising regional volumes to unit sum) and both gender and income inequality. We regress, within each 10y window, the female-male brain differences on a set of covariates including the socioeconomic variables as well as ICV differences and other confounds. **A**. Comparison of regression models including GII vs. regression models that do not include GII; the difference is significant (cluster-based permutation testing) from 28y onwards. **B**. Regression coefficients of models including GII; the curve of GII’s regression coefficients is regressed on age to highlight an increasing (possibly accumulating) effect throughout age. **C**. Including the countries’ Gross Domestic Product (GDP) in the regression models does not nullify the effect for GII but reduces it (we focus on a subsample where the GII and GDP do not correlate; see Fig SI-2A and Materials and Methods). **D**. Introducing the Gini index as a covariate, however, almost entirely suppresses the effect for GII in subsamples where the Gini index and GII are uncorrelated (Fig SI-2B).

### Grey matter differences linked to gender inequality reflect underlying economic disparities

As mentioned above, the GII is an aggregation of different dimensions, including economic but also other factors. We here question the weight of the economic dimension on the identified female-male differences. For this, we consider the countries’ GDP, a general measure of economic size; and the Gini index, which measures economic inequality. Intuitively, we can think of the GDP as a proxy of the average wealth of a country, and of the Gini index as a proxy of the variability of wealth across people.

As shown in **Fig 1A**, the GII is highly correlated to both the GDP (negatively) and the Gini factor (positively). These high correlations hamper the interpretation of the regression coefficients when including these as additional covariates (Mason & Perreault Jr., 1991). To be able to interpret the regression coefficients, we designed an ad-hoc constrained random sampling approach where, for each age window, we subsampled a subset of 5,000 subject pairs such that the GII and the Gini index/GDP are as non-correlated as possible across these pairs (see **Materials and Methods**). This way, we could run regressions on these samples where the regression coefficients can be interpreted. **Fig SI-2** shows the performance of the sampling algorithm in terms of the resulting correlation of the subsample (red), together with the correlation from the whole sample (blue). Superimposed on the panels, there are the number of pairs to subsample from within each window (the larger this number, the easier it is to achieve a low correlation). As observed, the (absolute) correlation is greatly reduced in the subsample, with windows up until 50y yielding a near-zero correlation between the GII and both the GDP and the Gini index.

Now, using these subsamples, we ran regressions where we included GDP and the Gini index as covariates together with the GII. Again, we repeated the analysis five times. The motivation to analyse the GDP together with the GII, given their large statistical association, is to rule out that the entirety of the effect is due to having generally better economic conditions, a factor that is known to shape the brain (Farah, 2017). **Fig 3C** shows that, when we include the GDP, the effect of the GII is reduced but remains broadly significant (p<0.01, cluster-based permutation testing). On the other hand, since the GII and the Gini index overlap in their definitions, analysing the GII and the Gini index together allows us to dissect which aspects of the GII may drive the observed female-male differences. **Fig 3D** shows that, when we add the Gini index as a covariate instead of the GDP, the effect of the GII is almost entirely suppressed, while the Gini index exhibits a substantially stronger effect for the whole examined age range. (Note that Fig 3B differs from Fig 3CD from the fact that these are computed on smaller subsamples).

Overall, this result suggests that most of the information in the GII is already contained in the Gini index, emphasising the impact of economic inequality between the sexes.

## Discussion

Economic and gender inequality has been shown to relate to brain differences amongst older adults (Legaz, Altschuler, …, Bruce, & Ibañez, 2024) as well as younger people (Zugman, et al., 2023). However, no study has examined, on a single data set and analysis, how gender inequality can affect our brains through the entire lifespan in a way that extends to diverse world populations. To have a more inclusive, wider-range perspective, we here exploit for the first time the vol2Brain dataset, which contains thousands of subjects from all over the world. Leveraging on this resource, we show that gender inequality is a significant explainer of brain differences in grey matter between females and males beyond brain size; that this effect may accumulate through life; and that the effect is (at least at an aggregated level, and per current data) directly or indirectly dominated by economic factors.

In the literature, it is disputed whether substantial and replicable differences between female and male brains exist that are not downstream consequences of trivial brain size differences (Eliot, Ahmed, Khan, & Patel, 2021). Here, we took care to separate brain size from *relative* size differences across regions, which is the focus of this paper. We did this in three ways, (i) during the vol2Brain core preprocessing, where we use a standardized advanced MRI preprocessing pipeline (denoising, bias field correction, intensity normalization) to minimise segmentation errors; (ii) by scaling all the regions’ volumes (in cm^3^) to sum up to 1.0, such that all subjects have the same total size regardless of their sex; and (iii) by explicitly correcting for ICV in both the out-of-sample predictions (regressing it out) and in the statistical tests (including it as an additional covariate). By doing this, we saw that GII is linked to relative brain differences in grey matter in a way that largely goes beyond brain size. But we also saw that, even after scaling the brain regions, ICV remains a significant covariate in our analysis, highlighting the multifaceted, complex influence of brain size on other brain measures (Wang, Hill-Jarrett, Buto, …, & Ackley, 2024).

Our data set has limited personal information (only sex, age and country), so we relied on country-level, structural socioeconomic variables. Although our predictions are done at the subject (or female-male-pair) level, this effectively means that the reported effects are strictly not based on individual but on country-average differences. Other data sets like the UK Biobank, whilst less inclusive in terms of their demography and age range, could be used to complement this analysis by incorporating subject-level socioeconomic information. While previous work has found tangible links between socioeconomic status and brain anatomy across subjects (Hyeokmoon, et al., 2022; Farah, 2017), how to distil measures of gender inequality at the individual level from this information is nonetheless difficult.

Cross-national neuroimaging datasets reflect not only population-level exposures but also structural biases in who gets scanned. In high-income countries, large volunteer-based cohorts typically underrepresent individuals with lower socioeconomic status or poorer health (Fry et al., 2017). In low- and middle-income countries, limited MRI access often confines scanning to private or clinical settings, skewing samples toward urban, wealthier, or symptomatic subgroups (World Health Organization, 2025; Jalloul et al., 2023). These sampling constraints limit both representativeness and generalizability, as models predicting cognitive traits or disease often fail on underrepresented profiles (Greene et al., 2022). Our results reveal that brain variation across a globally diverse sample tracks social inequality, underscoring the need to examine how context shapes both neural diversity and the framework we use to interpret it. Capturing these dynamics is critical for building predictive models that are both robust and biologically meaningful.

Driven by their need to be comparable between countries, the measures for gender and economic inequality used in this study also face limitations. For example, the GII measures female empowerment through national parliamentary representation, disregarding participation at the local government level due to lack of data in some countries. Similarly, the labour market dimension is incomplete due to missing information on incomes and unpaid work by women. Equally due to limited data availability, other dimensions related to both detrimental and protective gender-related factors of health (e.g., gender-based violence and social connectedness, respectively) are not captured through the GII.

In general, the total economical capacity of a country (measured by the GDP) is negatively correlated with both gender and income inequality, and these are positively correlated to each other. These high correlations hinder our capacity to distil explanatory factors with high granularity (Mason & Perreault Jr., 1991). However, these socioeconomic variables are not totally redundant, which is evidenced by those countries where these measures disagree (e.g. Iran or Greece). These countries were indeed the main drivers of our joint analysis of GII and GDP or the Gini index, given that our sampling approach draws heavily on them. We suggest these countries may be good targets to investigate this question further, possibly through dedicated data collection.

It should also be taken into account that processes like pregnancy can produce lasting changes in brain structure; and both have been associated with grey matter reductions (Hoekzema et al., 2017; Martínez-García et al., 2021; Servin-Barthet et al., 2025) — although the permanence of pregnancy-related changes remains uncertain (Luders et al., 2020). These transitions are not evenly distributed: in lower-income settings, structural inequality is associated with earlier pregnancies, higher fertility, and greater cumulative risk (cognitive, clinical, and social) even among individuals without a diagnosed pathology (Moguilner et al., 2024; Mukadam et al., 2019; Santamaria-Garcia et al., 2023; Zugman et al., 2023). Although GII does not measure biological transitions like pregnancy directly, its reproductive health dimension includes adolescent birth rate, which reflects the social conditions that influence the prevalence and timing of early pregnancies. Fertility rates are also correlate with the Gini index, suggesting that exposure to sex-specific biological processes such as pregnancy may vary systematically across countries (Perotti, 1996; Skakkebaek et al., 2022; Murray et al., 2018; Vollset et al., 2020). It remains crucial to establish whether the observed brain differences reflect disparities in life conditions or true pathological processes.

Overall, it is difficult to specifically pinpoint causal chains with observational data alone (Pearl, 2010), especially given that the non-imaging variables were not at the subject level, and given that the results primarily draw from aggregated population-level statistics of the neuroimaging data. However, our results suggest that the economic factor is indeed important. The fact that adding GDP as a covariate did not completely cancel the partial effect of GII, but adding the Gini index did, confirms that, even though the Gini index is not specific to gender inequality, it does reflect it as well. But the causal mechanisms by which this happens can be multifarious, for instance by acting as a mediator to other factors. Some of these factors may not be related to gender inequality in a direct way, for instance in a hypothetical scenario where higher socioeconomic inequality leads to more exposure to environmental toxins to which women are somehow more vulnerable.

A side conclusion of this work is about aggregating age groups together into joint analyses that control for age simply as a linear confound (Griffin, Gohil, Woolrich, Smith, & Vidaurre, 2025). Indeed, many studies of the relationship between inequality and socioeconomic status on the one hand and individual brain differences on the other hand do so by aggregating relatively large age groups. We here show that the nature of these relations can change throughout the life span, interacting with sex-related processes that, however shaped by social context, may be normative biological transitions.

Although we were careful to remove subjects that were identified to be part of large data collection initiatives (such as the Alzheimer’s Disease Neuroimaging Initiative or the Human Connection Project datasets), a standing limitation of the vol2Brain dataset (and of our study) is that there is no absolute guarantee that the origin and/or country of residence of the subject matches the country that was introduced in the online form. For example, a researcher in Australia may have collected data from some subjects of American origin and have introduced Australia as the country because this is where the research is based. Since there is no control for IP address, there is also a certain suspicion that some users introduced random countries; for example, there is an unusual number of subjects supposedly from North Korea, which we discarded. Overall, this represents a case of label noise. However, this does not affect our conclusions, since the consequence of label noise is, except in some cases, to reduce the prediction accuracy and not to inflate the observed effects. Therefore, our results stand not because of this issue, but in spite of it.

In summary, we provide descriptive evidence of at least some differences in the female and male brains that may not be “biologically programmed” but environmentally induced. Unlike socioeconomic measures, where the extent to which brain differences are a cause (Jensen, 1967) or a consequence (Gould, 1996) is controversial, it is highly unlikely that any hypothetical genetic factors underlying sex-related brain differences play any role in causing greater gender inequality within a country. Looking ahead, this work can serve as a foundation for future research aimed at disentangling how the complex, multivariate factors contributing to gender inequality may affect the brain across the lifespan.

## Materials and Methods

### Data and preprocessing

We included data acquired through the volBrain platform (Manjón & Coupé, 2016), an online system^3^ designed to automate the processing and quantitative analysis of brain T1-weighted MRI scans. The vol2Brain pipeline (Manjón, et al., 2022) is a robust algorithm designed to automatically segment T1 MR images into 135 different cortical and subcortical regions. The system streamlines image denoising, bias field correction, brain extraction, and tissue segmentation, providing cerebro-spinal fluid, white and grey matter volume measurements of different brain structures, which we retained for our analyses. It uses a multi-atlas label fusion approach based on manually labelled T1-weighted brain MRIs to perform its parcellation. Although the data have not yet been used, the pipeline has been successfully benchmarked to characterise the lifespan on separate data (Coupé, Catheline, Lanuza, Manjón, & Alzheimer’s Disease Neuroimaging Initiative, 2017). As well as optionally specifying country, sex and age, the users are given the option to let their data be used for research. We analysed only the subjects that gave consent and specified all demographic information, while also discarding scans that we detected to pertain to other mainstream data sets, such as the Alzheimer’s Disease Neuroimaging Initiative (ADNI) dataset, the Human Connection Project, and other initiatives —who may not necessarily be from the country from where the analysis was requested. Subjects submitted by any member of the research team were also discarded. This left us with 13,277 subjects in total from the total of 60.316 subjects analysed with this pipeline, covering 52 countries (39 countries with more than 10 subjects). While all the preprocessing had already been performed within the vol2Brain pipeline, here we took the additional step of scaling the grey matter volumes so that total sum is 1.0 for all subjects. This was intended to focus on relative brain differences and make our predictions more independent of ICV.

### Sex prediction

For the out-of-sample prediction of sex, we took 10y-long sliding windows from 10y- to 80y-old (±5y), implementing a stratified cross-validation scheme within each window. In detail, the brain data was standardized within the training set, and the training mean/s.d. was applied on the testing set such that it was approximately standardised with no data leakage. Also, within training, we regressed the following confounds out of the brain data: age (within the window span), the estimated SNR of the image, and the ICV. SNR and ICV were quantile-normalised using the Gaussian distribution as a reference to deal with occasional (missestimated) extreme values. This procedure yielded a matrix of deconfounding regression coefficients that were subsequently applied on the test set, as recommended by (Snoek, Miletić, & Scholte, 2019). We then performed sex classification using a classifier based on L_2_-regularised regression, choosing the regularisation parameter in a nested cross-validation loop (we did not use logistic regression due to excessive computational cost, given the large number of classifiers to fit, but the results were tested to be comparable). The entire procedure was embedded on a bootstrapping approach, where, for each age window and bootstrap repetition, we sampled 1000 subjects with a balanced distribution of sex, and ensuring no data leakage between train and test. This way, the results are comparable across age windows, which otherwise would have different numbers of subjects and sex proportions. We performed 100 repetitions of this bootstrapping procedure, which were used to compute a 95% confidence interval (DiCiccio & Efron, 1996). We used standard regression of this curve on age, and a t-test to assess the statistical significance of the slope.

### The sex-prediction generalisation matrix

To assess the generalisability across the lifespan of the brain features that allow us to predict sex, we devised a new metric: the age-prediction generalisation matrix. Akin to the temporal generalisation matrix in decoding analyses as used in cognitive science (Vidaurre, Cichy, & Woolrich, 2021), this simply consists of training one prediction model per window and testing on each other window, while ensuring the training and the testing sets do not overlap in the subjects they contain. This yields a (no. windows by no. windows) generalisation matrix, where the off-diagonal shows how well prediction models generalize to ages different to those on which they were trained. This was embedded in a bootstrapping procedure as before.

### Brain differences vs. structural inequality

For statistical testing of the relation between brain differences and structural inequality, we performed the analysis over 10y-long sliding windows (similarly to the prediction of sex). Within each age window, and for each country, we took each possible female-male pair within the country and computed the Euclidean distance of the (scaled) brain data between the male and the female. This gave us a distribution of male-female differences per country. We also gathered the exact age difference between the female-male pair of subjects, the difference in ICV, and the difference in SNR (where ICV and SNR were quantile normalised as before). Note that we include these as covariates in contrast with regressing them out beforehand, as we did for the out-of-sample prediction. Then, using multivariate regression, we regressed the pairwise brain distances on the age/ICV/SNR differences, including also the GII as a regressor (one value per pair, equal to the GII of the country that the pair belongs to). We compared the explained variance of this model with a regression model that did not include the GII as a regressor. Statistical significance was assessed using cluster-based permutation testing (Maris & Oostenveld, 2007), with the F-statistic as the base statistic. Additionally, using the same cluster-based permutation testing scheme, we assessed the statistical significance of each of the regressors with the t-statistic as the base statistic. We tested the slope of the GII covariate across the age span by regressing these curves on age, using a t-test. We also considered models with two structural inequality measures as regressors (e.g. GII and the Gini index, or GII and the GDP).

### Tackling covariate collinearity

To interpret the significance of the regressors, it is a problem when two of the covariates are highly collinear (Mason & Perreault Jr., 1991). This was the case of the GII, the Gini index, and the GDP. To tackle this issue, we devised an ad-hoc constrained random sampling approach based on a heuristic local search and simulated annealing (Bertsimas & Tsitsiklis, 1993), where for each regression we sampled 5,000 subject pairs such that the correlation (across pairs) between the two structural inequality metrics included in the model is (in absolute value) as low as possible. The solution of this optimisation problem is represented by a vector *x* with one element per country, such that the total sum is 5,000 and the maximum values of the vector elements correspond to the total number of pairs per country. Given some *x*, we can calculate the correlation between the two structural inequality metrics across the selected pairs in a computationally efficient way by using a modified version of the Pearson’s correlation equation:

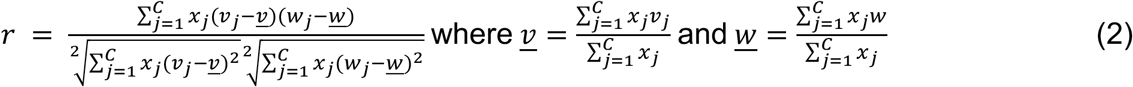

where *v* and *w* contain the values of the two structural inequality metrics for each country, and *C* is the number of countries.

The proposed algorithm minimizes this metric as follows: We start with a random selection of pairs *x* and try to improve the overall correlation by iteratively transferring pairs between the coordinates of *x*. At each iteration, we randomly choose two countries, such that one will transfer pairs to the other. The maximum number of pairs to transfer depends on a temperature parameter. How many pairs are proposed to be transferred is a number sampled from a discrete uniform distribution between 1.0 and a maximum number that depends on the temperature parameter. The temperature descends throughout the iterations given some “cooling” hyperparameter (Bertsimas & Tsitsiklis, 1993). If the transfer leads to a descent in absolute correlation (as per the equation above), it is accepted; if not, it is accepted with a probability that depends on both the increase in correlation and the same temperature parameter. This entire procedure is repeated ten times, from which we select the best solution (i.e. the one with the absolute lowest correlation). Fig **SI-2** illustrates the performance of the method.

## Acknowledgements

DV is supported by a Novo Nordisk Foundation Emerging Investigator Fellowship (NNF19OC-0054895), an ERC Starting Grant (ERC-StG-2019-850404) and by a DFF1 project from the Independent Research Fund Denmark (2034-00054B). This research was funded in part by the Wellcome Trust (215573/Z/19/Z). For the purpose of Open Access, the author has applied a CC BY public copyright licence to any Author Accepted Manuscript version arising from this submission.

## Supplemental Information

**Fig SI-1.**
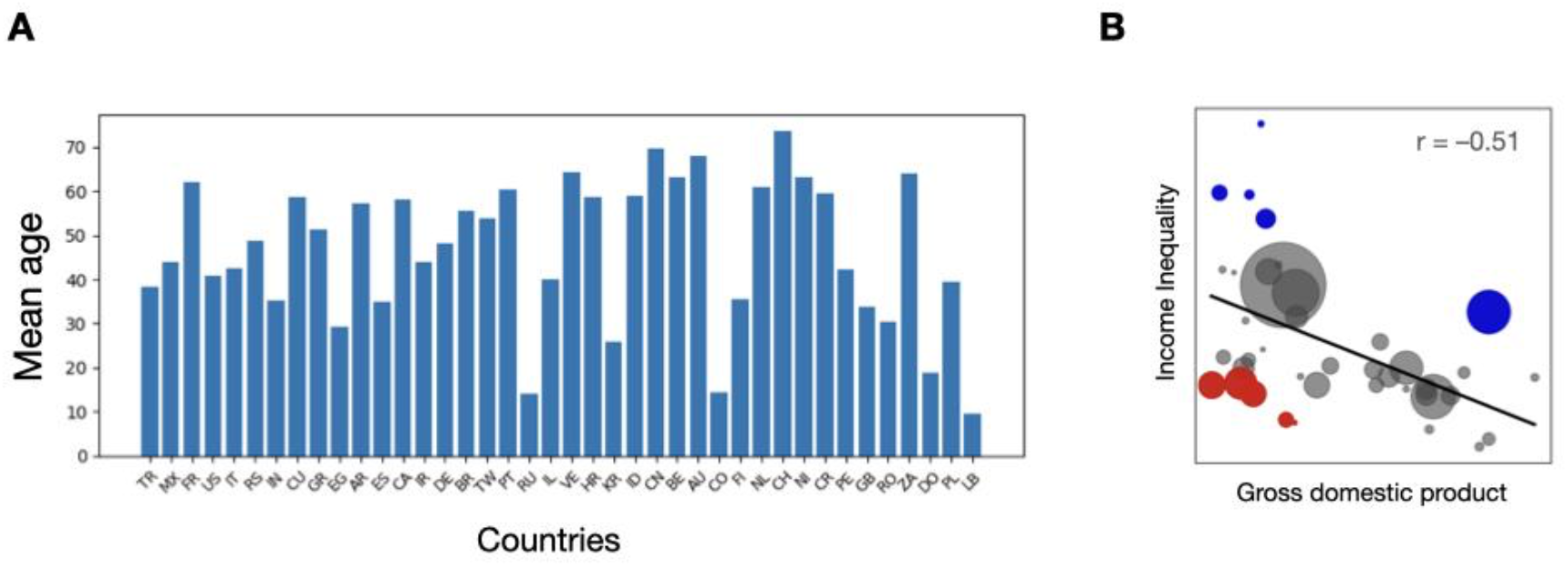
Further demographic information of the vol2Brain data set. **A**. Mean age per country (same order of countries as Fig 1). **B**. Income inequality versus GDP. Blue represents countries with relatively worse income inequality (Brazil, Colombia, EEUU, Venezuela, S. Africa), and red represents countries with relatively weaker GDP (Egypt, Croatia, India, Poland, Serbia).

**Fig SI-2.**
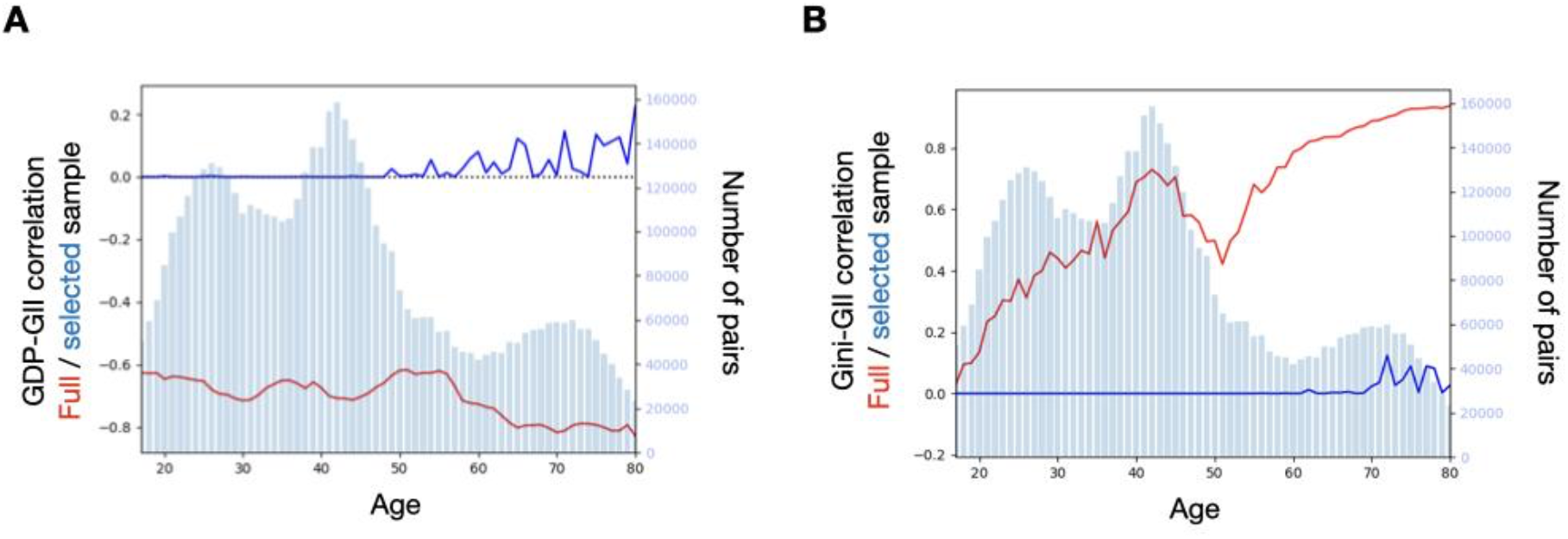
Assessment of the performance of our constrained random sampling approach across age groups, in terms of the correlation across subject pairs between the GII and two economy-related metrics. **A**. GII vs. GDP. **B**. GII vs. Gini index.

## Footnotes

1 https://hdr.undp.org/data-center/thematic-composite-indices/gender-inequality-index#/indicies/GII

2 https://data.worldbank.org

3 www.volBrain.net

## Bibliography

Arnold, A., & Gorski, R. (1984). Gonadal steroid induction of structural sex differences in the central nervous system. Annual review of neuroscience, 7, 413–442.

Baez, S., Castro-Aldrete, L., Britton, G., Ibañez, A., & Santuccione-Chadha, A. (2024). Enhancing brain health in the global south through a sex and gender lens. Nature Mental Health, 1–10.

Bertsimas, D., & Tsitsiklis, J. (1993). Simulated annealing. Statistical science, 8, 10–15.

Ceriani, L., & Verme, P. (2012). The origins of the Gini index: extracts from Variabilità e Mutabilità (1912) by Corrado Gini. The Journal of Economic Inequality, 10, 421–443.

Chekroud, A., Ward, E., Rosenberg, M., & Holmes, A. (2016). Patterns in the human brain mosaic discriminate males from females. Proceedings of the National Academy of Sciences, 14, E1968–E1968.

Coupé, P., Catheline, G., Lanuza, E., Manjón, J., & Alzheimer’s Disease Neuroimaging Initiative. (2017). Towards a unified analysis of brain maturation and aging across the entire lifespan: A MRI analysis. Human brain mapping, 38, 5501–5518.

De Vries, G., Rissman, E., Simerly, R., Yang, L.-Y., Scordalakes, E., Auger, C., … Arnold, A. (2002). A model system for study of sex chromosome effects on sexually dimorphic neural and behavioral traits. Journal of Neuroscience, 22, 9005–9014.

DiCiccio, T., & Efron, B. (1996). Bootstrap confidence intervals. Statistical science, 11, 189–228.

Eliot, L., Ahmed, A., Khan, H., & Patel, J. (2021). Dump the “dimorphism”: Comprehensive synthesis of human brain studies reveals few male-female differences beyond size. Neuroscience & Biobehavioral Reviews, 667–697.

Farah, M. (2017). The Neuroscience of Socioeconomic Status: Correlates, Causes, and Consequences. Neuron, 96, 56–71.

Gould, S. (1996). The Mismeasure of Man. WW Norton & company.

Griffin, B., Gohil, C., Woolrich, M., Smith, S., & Vidaurre, D. (2025). Does age moderate the relationship between brain structure and cognition? bioXriv.

Guiso, L., Monte, F., Sapienza, P., & Zingales, L. (2008). Culture, gender, and math. Science, 320, 1164–1165.

Hatzenbuehler, M., McLaughlin, K., Weissman, D., & Cikara, M. (2024). A research agenda for understanding how social inequality is linked to brain structure and function. Nature human behaviour, 8, 20–31.

Heise, L., Greene, M., Opper, N., Stavropoulou, M., Harper, C., Nascimento, M., & Zewdie, D. (2019). Gender inequality and restrictive gender norms: framing the challenges to health. The Lancet, 393, 2440–2454.

Hyeokmoon, K., Aydogan, G., Dagher, A., Bzdok, D., Christian C. R.,, Nave, G., … Koellinger, P. (2022). Human brain anatomy reflects separable genetic and environmental components of socioeconomic status. Science advances, 8, eabm2923.

Jensen, A. (1967). How much can we boost IQ and scholastic achievement? Harvard Educational Review, 39, 1–123.

Joel, D., Berman, Z., Tavor, I., Wexler, N., Gaber, O., Stein, Y., … Assaf, Y. (2015). Sex beyond the genitalia: The human brain mosaic. Proceedings of the National Academy of Sciences(112), 15468–15473.

Larraz, B. (2015). Decomposing the Gini Inequality Index: An Expanded Solution With Survey Data Applied to Analyze Gender Income Inequality. Sociological Methods & Research, 44, 508–533.

Legaz, A., Altschuler, F., …, Bruce, M., & Ibañez, A. (2024). Structural inequality linked to brain volume and network dynamics in aging and dementia across the Americas. Nature Aging, 1–16.

Lundberg, S., & Su-In, L. (2017). A Unified Approach to Interpreting Model Predictions. Advances in Neural Information Processing Systems, (pp. 4768–4777).

Mallard, T., Liu, S., Seidlitz, J., Ma, Z., Moraczewski, D., Thomas, A., & Raznahan, A. (2021). X-chromosome influences on neuroanatomical variation in humans. Nature Neuroscience, 24, 1216–1224.

Manjón, J., & Coupé, P. (2016). volBrain: an online MRI brain volumetry system. Frontiers in neuroinformatics, 10, 30.

Manjón, J., Romero, J., Vivo-Hernando, R., Rubio, G., Aparici, F., de la Iglesia-Vaya, M., & Coupé, P. (2022). vol2Brain: a new online pipeline for whole brain MRI analysis. Frontiers in neuroinformatics, 16, 862805.

Maris, E., & Robert Oostenveld, R. (2007). Nonparametric statistical testing of EEG-and MEG-data. Journal of Neuroscience Methods, 164, 177–190.

Mason, C., & Perreault Jr., W. (1991). Collinearity, power, and interpretation of multiple regression analysis. Journal of marketing research, 28, 268–280.

Mauvais-Jarvis, F., Merz, N., Barnes, P., …, Sandberg, K., & Suzuki, A. (2020). Sex and gender: modifiers of health, disease, and medicine. The Lancet, 396, 565–582.

Pearl, J. (2010). The foundations of causal inference. Sociological Methodology, 40, 75–149.

Ruigrok, A., Salimi-Khorshidi, G., Lai, M.-C., Baron-Cohen, S., Lombardo, M., Tait, R., & Suckling, J. (2014). A meta-analysis of sex differences in human brain structure. Neuroscience & Biobehavioral Reviews, 39, 34–50.

Snoek, L., Miletić, S., & Scholte, H. (2019). How to control for confounds in decoding analyses of neuroimaging data. Neuroimage, 184, 741–760.

Vidaurre, D., Cichy, R., & Woolrich, M. (2021). Dissociable Components of Information Encoding in Human Perception. Cerebral Cortex, 12, 5664–5675.

Wang, J., Hill-Jarrett, T., Buto, P., …, & Ackley, S. (2024). Comparison of approaches to control for intracranial volume in research on the association of brain volumes with cognitive outcomes. Human Brain Mapping, 45, e26633.

Young, W., Goy, R., & Phoenix, C. (1964). Hormones and Sexual Behavior: Broad relationships exist between the gonadal hormones and behavior. Science, 143, 212–218.

Zugman, A., Alliende, L., Medel, V., …, Sara, E.-L., & Crossley, N. (2023). Country-level gender inequality is associated with structural differences in the brains of women and men. Proceedings of the National Academy of Sciences, 20, e2218782120.

